# Microfluidic Agarose *µ*-Droplets for DNA-Encoded Chemical Library Screening

**DOI:** 10.64898/2026.02.15.706034

**Authors:** Yoojin Kim, Hayeon Kim, Jinhui Hong, Minseo Kang, Jaeyoung Bae, Sangyoon Ko, Minjae Kim, Byumseok Koh, Hakjoong Kim, Sang-Hee Shim, Kyubong Jo

## Abstract

DNA-encoded library (DEL) technology enables high-throughput small-molecule discovery but is typically performed using purified proteins under in vitro conditions that do not reflect native intracellular environments. Here, we present a microfluidic agarose μ-droplet platform for cellular-context DEL screening. The porous hydrogel droplets provide mechanically stable yet permeable microenvironments that protect weak protein–ligand interactions while enabling efficient buffer exchange and ligand diffusion. Importantly, mild cell permeabilization within droplets selectively retained chromatin-associated proteins, allowing screening directly in a cellular context. Using BRD4 as a model target, we validated intracellular ligand engagement by fluorescence imaging and super-resolution microscopy. Small-scale DEL screening selectively enriched JQ1 in both bead-based and cell-based formats, and large-scale DEL screening across millions of encoded compounds successfully identified hit molecules by sequencing. This agarose μ-droplet–based strategy expands DEL technology toward biologically relevant and chromatin-associated targets under near-native conditions.

## Introduction

DNA-encoded chemical libraries (DELs) have emerged as a powerful platform for high-throughput small-molecule discovery by coupling combinatorial chemistry with DNA barcoding and next-generation sequencing [1-3]. This approach enables the simultaneous screening of millions to billions of compounds in a single selection experiment and has led to the identification of numerous bioactive ligands against purified protein targets [2-4]. Despite these advances, conventional DEL screening is typically performed using isolated proteins immobilized on solid supports under simplified in vitro conditions [3,4].

Such formats do not fully capture native intracellular environments, where many target proteins function within multiprotein assemblies or chromatin-associated complexes [5]. In addition, the immobilization of purified proteins on solid supports, such as magnetic beads, may perturb higher-order structural organization or alter binding interfaces through surface constraints [6]. Protein conformation, ligand accessibility, and interaction dynamics can therefore differ substantially between purified and cellular contexts. As a result, conventional DEL methodologies may overlook ligands whose binding depends on higher-order structural organization or intracellular architecture [3].

To address this limitation, strategies that preserve target context while maintaining compatibility with DEL workflows are needed. Hydrogel-based microenvironments generated through microfluidics offer opportunities to provide mechanical stabilization, controlled molecular diffusion, and efficient solution exchange within defined compartments [7,8]. Agarose µ-droplets, in particular, form porous yet physically robust matrices that can confine biomolecules while permitting small-molecule access [8,9].

Here, we developed a microfluidic agarose µ-droplet platform for DEL screening that integrates both magnetic bead– based and cellular formats. Proteins were immobilized onto magnetic beads contained within agarose µ-droplets to maintain structural stability during selection, and this system was subsequently extended to cell-containing droplets under controlled permeabilization conditions. By combining droplet encapsulation, mild cell permeabilization, and chromatin retention, we established conditions that enable selective ligand enrichment against chromatin-associated targets [10]. Using BRD4 as a model system [11, 12], we validated intracellular ligand engagement by fluorescence and super-resolution microscopy [13] and demonstrated selective hit enrichment in both small- and large-scale DEL selections.

## Result

### Chromatin protein retention in agarose µ-droplets

To generate agarose µ-droplets encapsulating cells, we employed a flow-focusing microfluidic device (Fig. 1A) [14, 15]. The device consists of an aqueous inlet delivering cells suspended in molten low-gelling-temperature (LGT) agarose, and two opposing oil inlets that converge at a flow-focusing junction. As the aqueous stream passes through this junction, it is sheared by the continuous oil phase to form highly monodisperse µ-droplets [16]. A yellow heating plate placed beneath the mixing region (Fig. 1A) maintains the temperature to prevent premature gelation of agarose during droplet formation. Upon cooling, the agarose within the droplets so-lidifies to form a porous hydrogel matrix, and the droplets are collected in a hexagonal chamber.

**Fig. 1.**
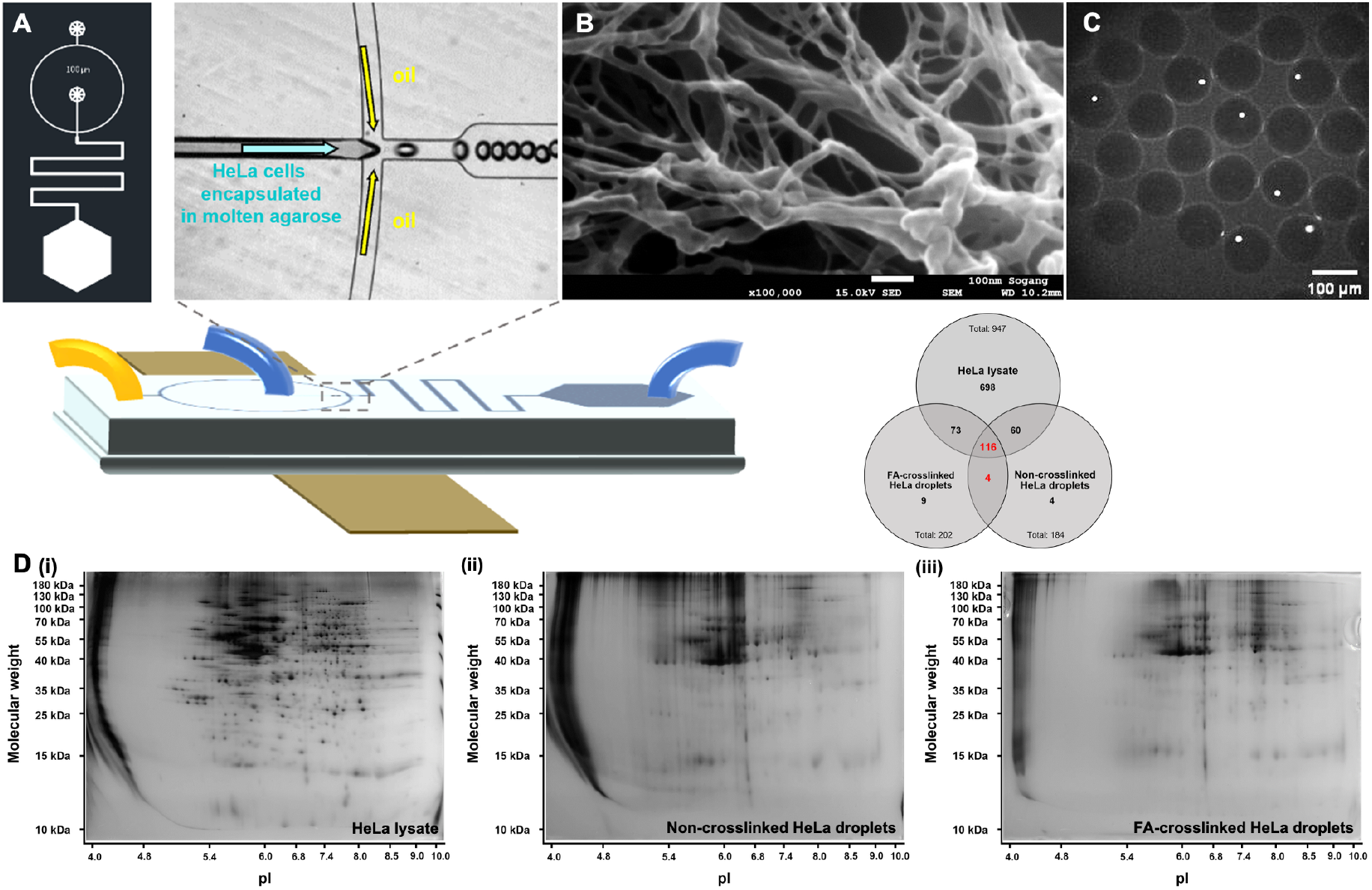
Design of a microfluidic device to generate agarose µ-droplets. **(A)** Schematic representation of the microfluidic flow-focusing device and the droplet generation region. An aqueous stream containing HeLa cells suspended in molten agarose is introduced through the central inlet and sheared by the continuous oil phase from opposing side channels at the flow-focusing junction, resulting in the formation of uniform agarose μ-droplets. **(B)** SEM image of the porous microstructure of 1.5% (w/w) low-gelling-temperature (LGT) agarose. Scale bar, 100nm. **(C)** Fluorescence microscope image of HeLa cells encapsulated within agarose μ-droplets stained with DAPI. **(D)** Two-dimensional gel electrophoresis (2DE) analysis of protein retention in agarose μ-droplet–encapsulated HeLa cells. Protein spot patterns are shown for (i) whole HeLa cell lysate, (ii) HeLa cells encapsulated in agarose μ-droplets and subjected to repeated washing under hypotonic conditions to induce cell lysis, and (iii) formaldehyde (FA)-treated HeLa cells encapsulated in agarose μ-droplets and processed under the same hypotonic lysis and washing conditions. Comparison of protein spot distributions reveals substantial retention of intracellular proteins after droplet encapsulation and washing, with significant overlap between washed droplets and FA-treated samples.

Scanning electron microscopy of 1.5% (w/w) LGT agarose revealed a highly porous and well-defined interconnected network structure (Fig. 1B), which provides mechanical stability while permitting free diffusion of small molecules through the matrix [17]. This porous architecture enables efficient molecular access to encapsulated components, allowing DNA-encoded compounds in solution to readily access target proteins. As a result, the agarose µ-droplets act as protective microenvironments that preserve the structural integrity of protein complexes throughout the screening process.

Fig. 1C shows a fluorescence microscopy image of generated agarose µ-droplets. In this experiment, HeLa cells were encapsulated within the droplets and stained with DAPI, and DAPI fluorescence was observed from the encapsulated HeLa cells, confirming successful single-cell encapsulation in an agarose µ-drolet.

This microfluidic method offers several advantages over conventional bulk emulsification. First, it produces uniform droplets with minimal polydispersity, ensuring consistent screening conditions [16,18]. Second, once the agarose droplets are solidified, the encapsulated protein–bead complexes are protected from shear forces of the solution, thereby preventing mechanical disruption of weak protein– ligand interactions. Third, the small size of the droplets (∼100 µm in diameter) allows rapid diffusion of small molecules into the porous hydrogel matrix, ensuring efficient access of DNA-encoded compounds to their target proteins. Fourth, the solidified droplets can be easily handled for solution exchange steps. The droplets settle naturally in test tubes under gravity, and their relatively large size allows for convenient buffer replacement without passing through filters. This feature facilitates repeated washing or solution exchange while minimizing physical perturbation, thereby preserving the integrity of protein– ligand complexes during the process.

Agarose µ-droplets were highly effective not only in trapping cells but also in retaining intracellular chromatin. To preserve chromatin within the droplets, cells needed to be lysed under conditions that avoided extensive protein denaturation. In this study, mild cell permeabilization was induced by treatment with 1× TE buffer (10 mM Tris, 1 mM EDTA, pH 8.0), which imposes hypotonic conditions that promote cellular swelling and partial membrane permeabilization [19]. Under such conditions, intracellular soluble proteins can be released from cytoplasm, while chromatin-associated components remain largely intact [20]. Because cells were physically confined within the agarose µ-droplets, most soluble cytoplasmic proteins diffused out through the porous droplet network and were efficiently removed during repeated washing steps. In contrast, chromatin-associated proteins remained bound to DNA and were selectively retained within the droplets [20]. The small size and porous structure of the agarose µ-droplets enabled rapid molecular diffusion and efficient solution exchange, allowing chromatin to remain trapped inside despite repeated, simple washing processes [21].

Hypotonic treatment induces pore formation in the cell and nuclear membranes, enabling DEL compounds to diffuse through the agarose matrix and access chromatin targets. Following mild cell permeabilization using 1× TE buffer, two-dimensional gel electrophoresis (2DE) was performed to evaluate whether chromatin-associated proteins are selectively retained within agarose µ-droplets. Three distinct samples were analyzed. Sample (i) consisted of untreated whole HeLa cells, which were directly lysed to obtain total cellular protein extracts. Sample (ii) comprised HeLa cells encapsulated in agarose µ-droplets, followed by TE buffer treatment and repeated washing to induce mild cell lysis and to remove soluble cytoplasmic proteins; the agarose droplets were subsequently dissolved, and the retained proteins were subjected to 2DE analysis. Sample (iii) was prepared in an analogous manner, except that HeLa cells were first treated with formaldehyde (FA) prior to droplet encapsulation.

In 2DE analysis, proteins are separated according to their isoelectric point and molecular weight, resulting in distinct protein spots [22]. 947 protein spots were observed in the whole HeLa cell lysate. Following cell permeabilization within agarose µ-droplets, 184 protein spots were observed without crosslinking, whereas FA crosslinking prior to permeabilization yielded 202 protein spots.

To further compare the protein populations preserved under different conditions, the overlap of protein spots identified by 2DE was analyzed. Both the number and distribution of protein spots were comparable between non-crosslinked and FA-crosslinked HeLa cell-containing µ-droplets following TE buffer treatment. This similarity suggests that chromatin-associated proteins remain bound to DNA under mild TE-induced lysis conditions, without the need for additional chemical crosslinking [20].

### Selective intracellular targeting in agarose µ-droplets

To establish that protein-ligand interactions can be specifically detected within agarose µ-droplets, we employed a fluorescent JQ1-DNA conjugate (JQ1-oligo-Cy5), hereafter referred to as the binding probe (BP), as a model probe targeting BRD4 (Fig. 2A). JQ1 is a well-characterized small-molecule ligand of BRD4 [23] and was therefore selected as a positive control to evaluate target engagement under droplet-confined conditions. Conjugation to a Cy5-labeled DNA oligonucleotide enables direct fluorescence-based visualization of probe localization within agarose µ-droplets. To validate whether the fluorescent probe could specifically report protein-ligand interactions within agarose µ-droplets, we next examined the binding behavior of BP in both bead-based and cell-based droplet systems after incubation with BP followed by washing steps to remove unbound probe (Fig. 2B). The images shown in Fig. 2B are overlays of bright-field and Cy5 fluorescence channels, enabling simultaneous visualization of the agarose µ-droplets, encapsulated magnetic beads or cells, and the localization of the fluorescent probe.

**Fig 2.**
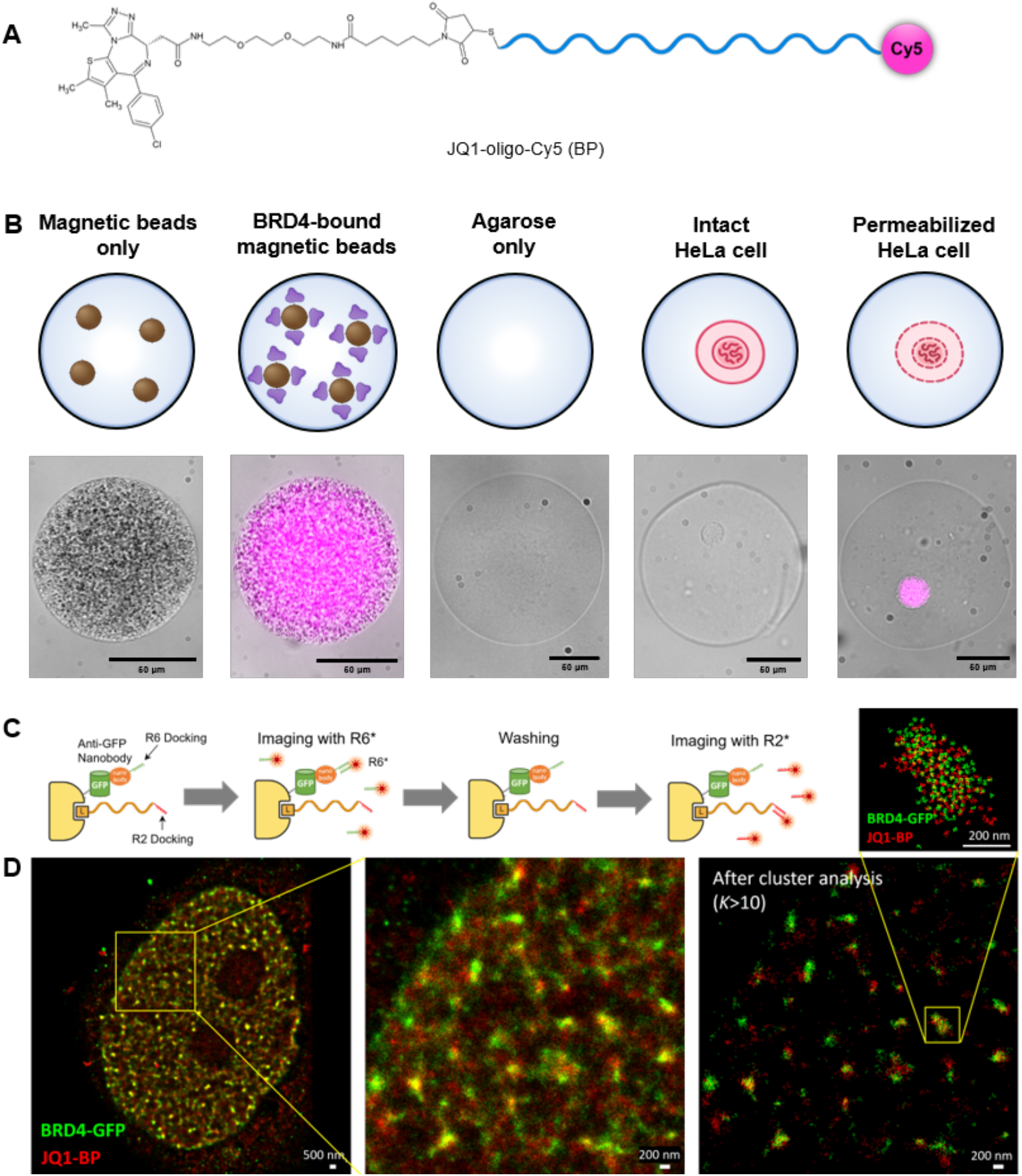
Validation of JQ1-oligo-Cy5 for specific detection of BRD4 in agarose μ-droplets. **(A)** Chemical structure of the JQ1-oligo-Cy5 conjugate used as a fluorescent probe (BP). JQ1, a selective BRD4 ligand, is covalently linked to a DNA oligonucleotide and a Cy5 fluorophore, allowing fluorescence-based detection of BRD4 binding. **(B)** Schematic illustrations (top) and corresponding fluorescence images (bottom) of agarose μ-droplets containing magnetic beads only, BRD4-bound magnetic beads, agarose alone, intact HeLa cells, or permeabilized HeLa cells after incubation with JQ1-oligo-Cy5. Strong Cy5 fluorescence was observed in droplets containing BRD4-bound magnetic beads and permeabilized HeLa cells, whereas no detectable Cy5 fluorescence was observed in magnetic beads only, intact HeLa cells, and agarose-only droplets. Scale bars, 50μm. **(C)** Schematic illustration of two-color Exchange-PAINT imaging using orthogonal DNA docking-imager strand pairs. An R6 docking strand conjugated to an anti-GFP nanobody was used to label GFP-BRD4, while an R2 docking strand was incorporated into the JQ1-based probe (JQ1-BP). Sequential super-resolution imaging was performed using R6* and R2* imager strands to independently localize BRD4 and bound JQ1, respectively. **(D)** Two-color Exchange-PAINT super-resolution images of GFP-BRD4-expressing Cos7 cells showing GFP-BRD4 (green) and the JQ1-based probe (JQ1-BP, red). Left, whole-nucleus view; middle, magnified view of the boxed region highlighting nanoscale clustering of BRD4 and JQ1. Right, the same region after Q-PAINT-based cluster filtering (K > 10), revealing higher-order nanoclusters containing both BRD4 and JQ1. Enrichment of JQ1-BP localizations at GFP-BRD4 clusters indicates that JQ1 preferentially localizes to BRD4-enriched nuclear regions.

As an initial validation step, BP binding was examined in bead-based agarose µ-droplets. Strong Cy5 fluorescence was observed in agarose µ-droplets containing BRD4-bound magnetic beads, with fluorescence signals colocalized with the bead regions, indicating specific association of BP with immobilized BRD4. In contrast, droplets containing magnetic beads alone showed no detectable Cy5 fluorescence, demonstrating that BP does not nonspecifically interact with the bead surface under these conditions. These observations indicate that BP can freely diffuse into the agarose µ-droplets to access immobilized BRD4 [24], and that unbound probe is effectively removed during subsequent washing steps. These results confirm that the observed fluorescence signals arise from specific BP-BRD4 interactions rather than nonspecific retention within the agarose matrix or on the magnetic bead surface. Thus, the agarose µ-droplet environment preserves protein accessibility while enabling selective detection of protein-ligand binding after removal of unbound probe.

We further examined whether BP could access intracellular targets in a cellular context. To this end, intact HeLa cells were encapsulated in agarose µ-droplets and subjected to post-encapsulation washing conditions that modulate cellular permeability, as established above. Briefly, PBS washing was used to maintain isotonic conditions and preserve intact cellular membranes, while TE washing was applied to induce mild permeabilization within droplets.

Whereas HeLa cell–containing droplets maintained under isotonic conditions showed no detectable Cy5 fluorescence, droplets subjected to TE washing exhibited a pronounced Cy5 signal specifically associated with the encapsulated cells. These observations indicate that intracellular access of BP is enabled only after permeabilization induced at the droplet stage, and that BP does not gain intracellular access in intact cells under isotonic conditions due to the impermeability of the plasma membrane. Collectively, these results establish that cellular permeability can be modulated by post-encapsulation washing conditions, enabling selective engagement of intracellular BRD4 in a droplet-confined environment.

Although fluorescence imaging demonstrated intracellular accessibility of BP, diffraction-limited microscopy could not resolve whether the observed signals represented direct ligand–target engagement or nonspecific probe accumulation. To directly verify nanoscale BRD4-JQ1 interactions in a cellular context, we next performed super-resolution imaging using DNA-PAINT (Points Accumulation for Imaging in Nanoscale Topography) [25].

To this end, we implemented a two-color Exchange-PAINT strategy based on orthogonal DNA docking-imager strand pairs (Fig. 2C) [25]. An R6 docking strand was conjugated to an anti-GFP nanobody to label BRD4-GFP, while an R2 docking strand was incorporated into the JQ1-based probe (JQ1-BP). Super-resolution imaging was performed sequentially to avoid cross-talk between channels. BRD4-GFP was first imaged using an R6-complementary fluorescent imager strand (R6*), enabling nanoscale localization of the target protein. After removal of the R6* imager strand by washing, an R2-complementary imager strand (R2*) was introduced to selectively visualize BP molecules bound to BRD4. Two-color Exchange-PAINT imaging of BRD4-GFP-expressing Cos7 cells revealed that both BRD4 and JQ1 signals were organized into discrete nanoscale clusters within the nucleus (Fig. 2D). Notably, JQ1-BP localizations detected by R2* imaging were highly enriched at BRD4-GFP clusters identified by R6* imaging, rather than being randomly distributed throughout the nucleus, providing direct nanoscale evidence of ligand engagement at BRD4-enriched domains.

At the single-molecule level, individual BRD4 and JQ1 molecules appeared as clusters of multiple localization events, consistent with the transient binding kinetics characteristic of DNA-PAINT [25]. Application of Q-PAINT-based cluster filtering (K > 10) [26] revealed that these molecular clusters further assembled into higher-order nanoclusters containing both BRD4 and JQ1. The persistence of BRD4-JQ1 co-clustering after stringent filtering indicates that the observed super-resolution signals reflect specific intracellular ligand-target interactions rather than coincidental spatial overlap or nonspecific probe retention.

Collectively, these results directly confirm that JQ1 engages BRD4 at the nanoscale within a cellular environment and validate that the fluorescence signals observed in agarose µ-droplet assays originate from genuine intracellular protein–ligand interactions.

### Small-scale proof of concept DEL screening

Prior to large-scale DEL screening, we first sought to validate the feasibility of our agarose µ-droplet based screening platform using a small, well-defined DNA-encoded library. To this end, a small-scale DEL consisting of four representative compounds was employed as a proof-of-principle library (Fig. 3A). This library was designed to include compounds with distinct and known binding characteristics, enabling straightforward evaluation of target specificity and background enrichment. The four-compound library comprised JQ1, a canonical BRD4 bromodomain inhibitor, two off-target binders (GL-CBS and MTX, targeting carbonic anhydrase II and dihydrofolate reductase, respectively [27, 28]), and benzoic acid as a non-binding negative control. Importantly, this small-scale DEL was synthesized externally by a collaborating institution and provided as a ready-to-use encoded library. By first applying this minimal library, we aimed to confirm that target-dependent enrichment and signal discrimination could be reliably achieved within agarose µ-droplets before proceeding to more complex, large-scale DEL selections [29]. To examine whether the small-scale DEL could specifically report BRD4-dependent enrichment in a bead-based context, we next performed small-scale DEL screening in agarose µ-droplets containing magnetic beads under two conditions: droplets containing magnetic beads that were subsequently incubated with BRD4 and droplets containing magnetic beads alone as a background control (Fig. 3B).

**Fig 3.**
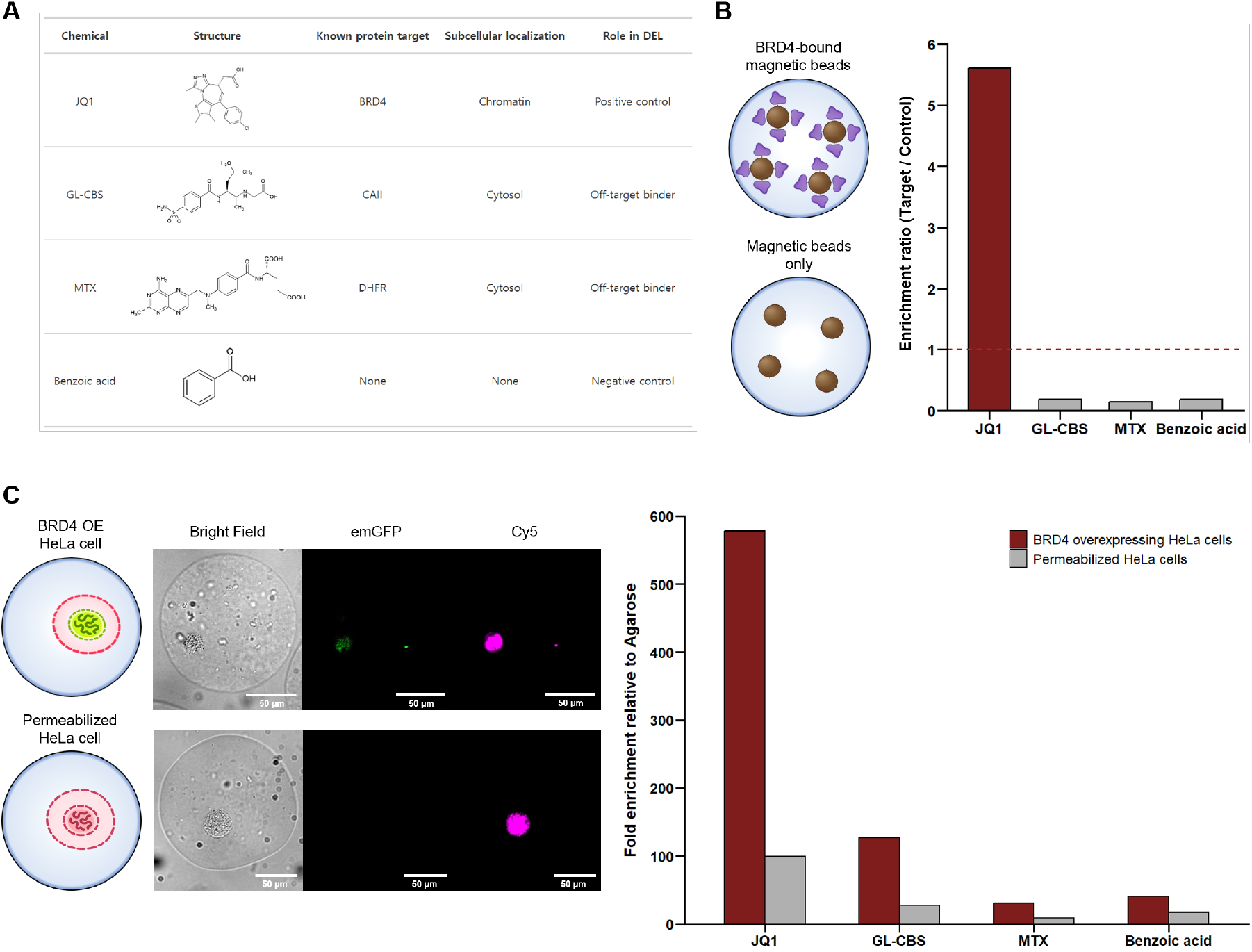
Proof-of-concept DEL screening against BRD4 using a small-scale four-compound library in agarose μ-droplets. **(A)** Composition of the four-compound library used for proof-of-concept DEL screening, including JQ1 as a positive control, GL-CBS and methotrexate (MTX) as off-target binders, and benzoic acid as a negative control. **(B)** Enrichment ratios (Target / Control) derived from nanopore sequencing data following small-scale DEL screening in agarose μ-droplets containing BRD4-bound magnetic beads (target) or magnetic beads only (control). JQ1 exhibited strong enrichment above the threshold (Target / Control = 1), whereas other compounds showed minimal enrichment. **(C)** Quantitative analysis of small-scale DEL screening performed in cell-containing agarose μ-droplets, in which DEL enrichment was quantified by qPCR following screening. Agarose-only μ-droplets were used as a control condition, while μ-droplets containing permeabilized HeLa cells or BRD4-overexpressing HeLa cells were treated as target conditions. JQ1 showed the highest enrichment among tested compounds and displayed a pronounced preference for BRD4-overexpressing HeLa cells compared to permeabilized cells. Representative bright field, emGFP, and Cy5 fluorescence images are shown on the right. Scale bars, 50μm. emGFP fluorescence was observed specifically in BRD4-overexpressing HeLa cells. As both HeLa cell conditions were permeabilized, Cy5 fluorescence originating from BP was detected in droplets containing either permeabilized or BRD4-overexpressing HeLa cells.

Following DEL incubation and washing within droplets, recovered DNA barcodes were subjected to nanopore sequencing [30], and enrichment ratios were calculated for each compound by comparing target and control nanopore sequencing read counts. Analysis of sequencing results revealed a pronounced and selective enrichment of the JQ1-encoded barcode in droplets containing BRD4-bound magnetic beads, whereas minimal enrichment was observed in bead-only control droplets. In contrast, GL-CBS, MTX, and benzoic acid showed enrichment ratios close to background levels, indicating negligible non-specific capture under these conditions. These data demonstrate that bead-based DEL screening within agarose µ-droplets, coupled with nanopore sequencing-based decoding, robustly identifies a known BRD4 binder while effectively suppressing off-target and background signals.

We next extended this proof-of-concept screening to a cell-based format by encapsulating HeLa cells in agarose µ-droplets. Three sample conditions were prepared to evaluate target-dependent enrichment within droplets. First, parental HeLa cells were encapsulated into agarose µ-droplets and subsequently rendered permeable using the droplet-based permeabilization strategy established above. Second, HeLa cells transiently transfected with a BRD4-emGFP expression plasmid were used to generate a BRD4-overexpressing model. These cells were similarly encapsulated into agarose µ-droplets and processed under the same permeabilization conditions. As a control, cell-free agarose µ-droplets were prepared and processed in parallel. Fluorescence imaging was performed to visualize intracellular target presence within HeLa-cell containing agarose µ-droplets. emGFP fluorescence was used to confirm BRD4 expression, while Cy5 fluorescence from BP was monitored to assess probe accessibility in permeabilized HeLa cell droplets. In BRD4-overexpressing HeLa cell droplets, emGFP fluorescence originating from GFP-BRD4 expression was observed together with Cy5 fluorescence arising from BP, confirming the presence of intracellular BRD4 and effective probe accessibility within droplets. In contrast, permeabilized parental HeLa cell droplets showed Cy5 fluorescence from BP but no detectable emGFP signal.

We then performed small-scale DEL screening in HeLa cell–containing agarose µ-droplets under three sample conditions: permeabilized parental HeLa cell droplets, permeabilized BRD4-overexpressing HeLa cell droplets, and cell-free agarose droplets as a control. Following DEL incubation and washing, recovered DNA barcodes corresponding to individual compounds were quantified by qPCR using compound-specific primers. Compound enrichment was calculated by determining ΔCt values between target droplets and agarose-only control droplets (ΔCt = Ct_control − Ct_target), and relative enrichment was expressed as 2^ΔCt [31].

qPCR analysis revealed that JQ1 exhibited the highest level of enrichment among the tested compounds in HeLa cell-containing droplets compared to the agarose control. Notably, JQ1 enrichment was markedly increased in BRD4-overexpressing HeLa cell droplets relative to parental HeLa cell droplets, whereas GL-CBS, MTX, and benzoic acid remained near background levels under both cellular conditions. These results indicate that JQ1 enrichment in the small-scale DEL is closely associated with the presence and abundance of intracellular BRD4.

Together, these results demonstrate that the agarose µ-droplet platform enables reliable discrimination of target-specific binders using both sequencing-based (nanopore) and PCR-based readouts. This validation at a small-scale establishes a foundation for subsequent large-scale DEL screening using the same platform.

### Large-scale DEL screening in agarose µ-droplets

To further validate the applicability of the droplet-based screening platform to large and diverse chemical spaces, we performed DEL screening using a multi-component library synthesized by the Korea Research Institute of Chemical Technology (KRICT). The library comprised three distinct core scaffolds—pyrimidine, trifunctional benzene, and chiral proline cores—each assembled from three building block positions, resulting in a 96 × 96 × 96 combinatorial space per core scaffold [32,33]. Screening was conducted across four sample conditions: magnetic bead–only droplets, BRD4-bound magnetic bead droplets, permeabilized HeLa cell droplets, and BRD4-overexpressing HeLa cell droplets. Following screening, library composition and read count distributions were assessed by nanopore sequencing [30], and read counts corresponding to each core scaffold were extracted for analysis. In this experimental design, magnetic bead–only droplets and permeabilized parental HeLa cell droplets served as control conditions, while BRD4-bound magnetic bead droplets and BRD4-overexpressing HeLa cell droplets were used as target-containing samples, respectively. Across both bead-based and cell-based screening formats, target-containing samples exhibited distinct read count distributions compared to control conditions. Notably, despite being introduced at equal proportions, the chiral proline core exhibited the highest read counts among the three core scaffolds. To visualize enrichment patterns at the level of individual building block combinations, three-dimensional plots were constructed for each core scaffold based on the top 200 BB1–BB2–BB3 combinations ranked by nanopore sequencing read count (Fig. 4B) [34], where the x, y, and z axes correspond to BB1, BB2, and BB3 positions, respectively. In these representations, each point denotes a unique BB1–BB2–BB3 combination, and color intensity reflects the corresponding read count. Notably, combinations exceeding a read count threshold of 65 were further emphasized by increasing marker size, facilitating rapid identification of highly enriched species within the vast combinatorial space. Collectively, these results demonstrate that the droplet-based DEL screening platform enables interrogation and visualization of large-scale libraries across distinct biochemical contexts.

**Fig. 4.**
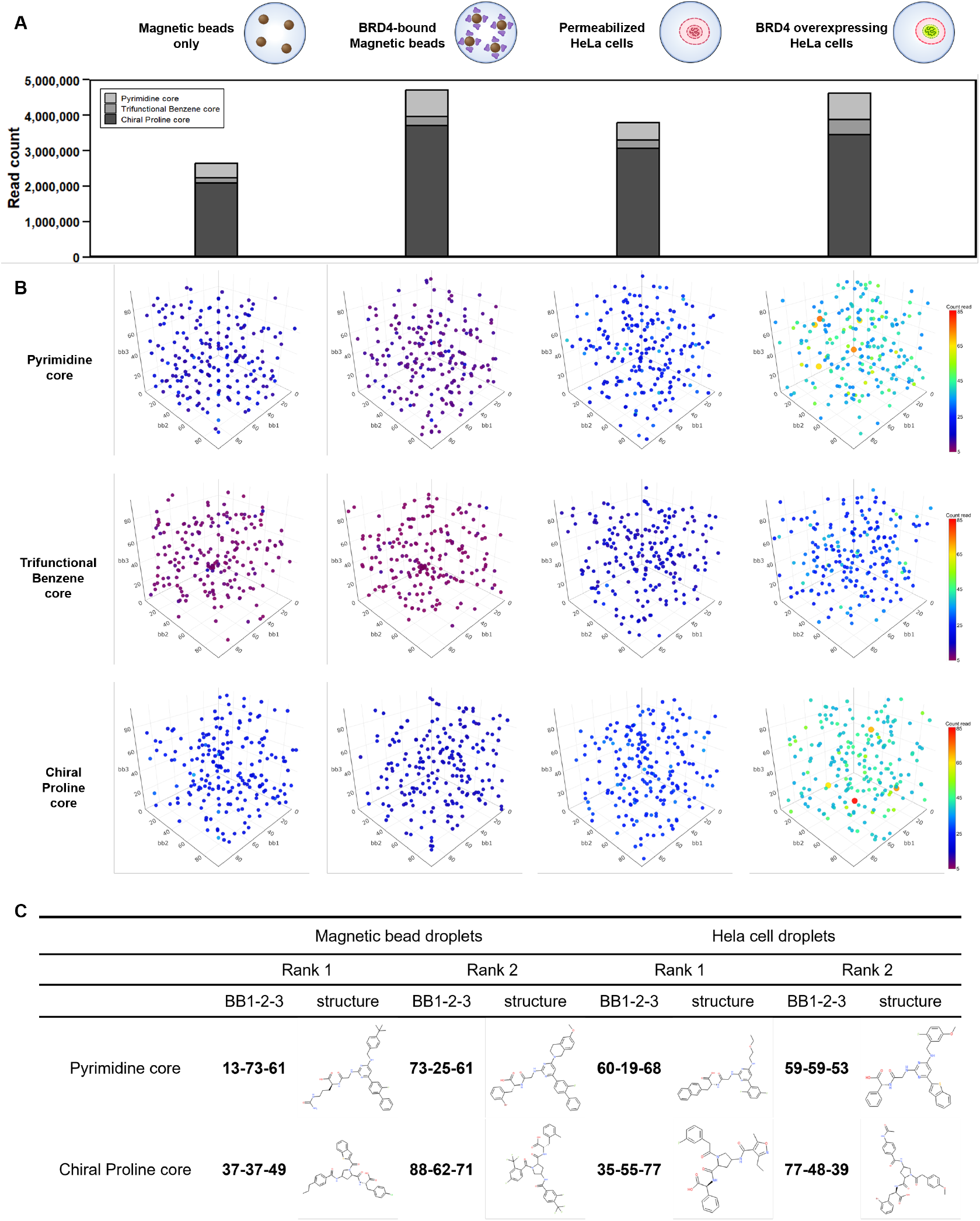
Large-scale DEL screening against BRD4 using a multi-component library in agarose μ-droplets. **(A)** Aggregated nanopore sequencing read count distributions for each core scaffold across four screening conditions: magnetic bead–only droplets, BRD4-bound magnetic bead droplets, permeabilized HeLa cell droplets, and BRD4-overexpressing HeLa cell droplets. The DNA-encoded library consisted of three distinct core scaffolds (pyrimidine, trifunctional benzene, and chiral proline cores), each comprising a 96 × 96 × 96 combinatorial spaces. **(B)** Three-dimensional enrichment maps for each core scaffold across the four screening conditions, displaying the top 200 BB1–BB2–BB3 combinations ranked by nanopore sequencing read count. Each axis (x, y, and z) corresponds to one building block position (BB1, BB2, and BB3), and individual points represent distinct BB1– BB2–BB3 combinations. Point color indicates read count abundance, while combinations exceeding a read count threshold of 65 are additionally highlighted by increased marker size to emphasize highly enriched species. **(C)** Chemical structures of the top enriched compounds identified from large-scale DEL screening. Enrichment was calculated as the ratio of nanopore sequencing read counts between target and control samples. For each core scaffold, the two compounds with the highest target-to-control enrichment are shown for both magnetic bead droplet and HeLa cell droplet screening conditions. The corresponding BB1-BB2-BB3-derived structures are displayed.

Based on nanopore sequencing read counts, compound-level enrichment was calculated as the ratio of nanopore read counts in target samples to those in corresponding control samples [29]. For each screening condition, compounds were ranked according to their target-to-control enrichment. Fig. 4C presents the top two enriched compounds for the pyrimidine and chiral proline cores under both bead-based and cell-based screening conditions, with the corresponding BB1-BB2-BB3 derived chemical structures shown. The trifunctional benzene core was excluded from compound-level enrichment analysis because read counts were low across both target and control samples, precluding statistically reliable enrichment calculations.

## Discussion

In this study, we established a microfluidic agarose µ-droplet platform that enables DEL screening under structurally preserved and context-aware conditions. Conventional DEL approaches are typically performed using purified proteins immobilized on solid supports in simplified in vitro environments. While effective for high-throughput ligand discovery, such formats often fail to recapitulate the structural and compositional complexity of intracellular protein assemblies. By integrating agarose µ-droplet encapsulation with controlled permeabilization strategies, our platform addresses this limitation and extends DEL screening into more biologically relevant contexts.

A central feature of this system is the use of LGT agarose to generate porous, mechanically stable droplets that confine cells or protein–bead complexes while permitting efficient diffusion of small molecules. The porous hydrogel matrix enables rapid buffer exchange and removal of unbound compounds, yet protects weak or transient protein–ligand interactions from mechanical disruption. This structural stabilization is particularly important for chromatin-associated proteins and multiprotein assemblies, where higher-order organization can strongly influence ligand accessibility and binding affinity. The observed retention of chromatin-associated proteins following hypotonic 1× TE-mediated permeabilization, without the need for chemical crosslinking, highlights the ability of agarose droplets to preserve functional macromolecular architectures during screening.

Using BRD4 as a model chromatin-associated target, we validated that protein–ligand interactions within agarose droplets can be detected with high specificity. Fluorescence-based assays demonstrated selective enrichment of JQ1 within both bead-based and cell-containing droplets under permeabilized conditions, while nonspecific retention was minimal. Importantly, two-color DNA-PAINT super-resolution imaging provided direct nanoscale evidence of intracellular BRD4–JQ1 co-localization. These imaging results confirm that the fluorescent signals observed in droplet assays reflect genuine molecular engagement rather than passive probe accumulation. The ability to correlate droplet-based ligand enrichment with independent super-resolution validation strengthens the mechanistic interpretation of the screening outcomes.

Proof-of-concept screening using a small-scale four-compound DEL further demonstrated that the platform enables reliable discrimination of target-specific binders. Both nanopore sequencing and qPCR-based readouts identified JQ1 as selectively enriched in BRD4-containing droplets, with negligible enrichment of off-target or non-binding controls. Notably, enrichment increased in BRD4-overexpressing HeLa cells relative to parental cells, indicating that ligand enrichment quantitatively reflects intracellular target abundance. These findings suggest that the droplet-based system preserves functional dependency between ligand selection and target expression level within a cellular context.

Large-scale screening of a multi-component DEL comprising nearly one million theoretical combinations per core scaffold further demonstrated the scalability of the platform. Distinct enrichment patterns were observed across bead-based and cell-based formats, indicating that ligand selection can vary depending on biochemical context. The differential read count distributions among core scaffolds suggest that scaffold architecture influences compatibility with preserved chromatin-associated environments. Importantly, the ability to visualize enrichment at the individual building block level using three-dimensional combinatorial mapping provides a powerful framework for decoding structure–enrichment relationships within complex libraries.

While the platform significantly advances context-preserving DEL screening, several considerations remain. Cellular permeabilization must be carefully controlled to balance intracellular accessibility with structural preservation. In addition, while hypotonic treatment retained chromatin-associated proteins effectively, other classes of targets may require optimized permeabilization strategies. Furthermore, although trifunctional benzene cores exhibited low read counts in the present study, future optimization of library representation or sequencing normalization may improve statistical robustness for such scaffolds.

Overall, the agarose µ-droplet system bridges the gap between simplified in vitro DEL screening and ligand discovery in cellular contexts. By integrating microfluidic encapsulation, hydrogel-based stabilization, super-resolution validation, and high-throughput sequencing, this platform enables context-preserving selection against chromatin-associated proteins and complexes. This approach may broaden small-molecule discovery toward previously inaccessible intracellular targets and higher-order macromolecular assemblies.

## Acknowledgements

This work was supported by the National Research Foundation of Korea (NRF) grant RS-2023-00258599. The authors thank the Korea Research Institute of Chemical Technology (KRICT) for providing the DNA-encoded chemical libraries used in this study.

## Author contributions

K.J and S.S conceived and designed the study. Y.K., H.K., J.H., M.K. and J.B. performed experiments and analyzed the data. S.K. and S.S performed DNA-PAINT imaging and analysis. M.K. and H.K. synthesized the small-scale DNA-encoded library. Y.K. wrote the manuscript. All authors approved the final manuscript.

## Materials and Methods

### Chemical and Materials

SU-8 2050 photoresist and SU-8 developer were obtained from Kayaku Advanced Materials (Westborough, MA, USA). Polydimethylsiloxane (PDMS) and curing agent were purchased from K1 Solution (Gwangmyeong, Korea). Ultra-low-gelling-temperature agarose (catalog no. A5030) and low-gelling-temperature agarose (catalog no. A4018) were obtained from Sigma-Aldrich (St. Louis, MO, USA). Tris(2-carboxyethyl)phosphine hydrochloride (TCEP) was obtained from Sigma-Aldrich (St. Louis, MO, USA). Thiol-modified and Cy5-labeled oligonucleotides were synthesized by Bioneer (Daejeon, Korea). The BRD4 bromodomain plasmid (pHis-BRD4 BD1, item #196544), GFP-BRD4 expression plasmid (GFP-BRD4, item #65378) was obtained from Addgene (Watertown, MA, USA). Escherichia coli strains DH5α and BL21 (DE3) were obtained from Enzynomics (Daejeon, Korea). Lipofectamine™ 3000 Transfection Reagent and Ni–NTA magnetic beads (HisPur™ Ni-NTA Magnetic Beads, catalog no. 88831) were purchased from Thermo Fisher Scientific (Waltham, MA, USA). Ni–NTA agarose resin and disposable gravity columns were purchased from Qiagen (Venlo, Netherlands). (+)-JQ1 maleimide (catalog no. 7576) was obtained from Tocris Bioscience (Bristol, UK), AccuPower® ProFi Taq PCR PreMix and Accu-Power® 2X GreenStar™ qPCR MasterMix were obtained from Bioneer (Daejeon, Korea). The ligation sequencing kit (SQK-LSK114) and R10.4.1 flow cells (FLO-MIN114) were purchased from Oxford Nanopore Technologies (Oxford, UK). Unless otherwise specified, all other chemicals were of analytical grade.

### Fabrication of PDMS Microfluidic Device

The mold for the droplet generator was fabricated using a standard soft lithography process. The procedure followed the datasheet protocol of SU-8 2050 (Kayaku Advanced Materials, USA). Silicon wafers were sequentially rinsed with distilled water, isopropyl alcohol, acetone, isopropyl alcohol, and distilled water, and then dried with nitrogen gas. The droplet generator patterns were defined using a chrome photomask (Advanced Micro Engineering & Design, Gunpo, Korea). A layer of SU-8 2050 photoresist was spin-coated onto the cleaned silicon wafer using a spin coater (SPIN-1200D, Midas System, Daejeon, Korea), followed by a soft bake. The coated wafer was then exposed to UV light using a mask aligner (MDA-400LJ, Midas System, Daejeon, Korea) through the photomask. After post-exposure baking, the substrate was developed with SU-8 developer and rinsed with isopropyl alcohol and distilled water to remove residual photoresist.

The PDMS microfluidic device was prepared by mixing SYLGARD™ 184 silicone elastomer base and curing agent (Dow Corning, USA) at a 10:1 (w/w) ratio. Prior to casting, short segments of tubing were cut and fixed onto the mold at the designated inlet and outlet positions using adhesive. The PDMS mixture was then poured over the mold, covering the embedded tubing segments, and cured in an oven at 65 °C for at least 4 h. After curing, the PDMS replica was peeled off from the mold. External tubing was connected to the inlet and outlet ports of the PDMS layer, which was subsequently bonded to a glass slide by oxygen plasma treatment (Cute Basic, Femto Science, Korea). After bonding, additional PDMS mixture was applied around the edges to securely fix the PDMS layer to the glass slide [35].

### Preparation of HeLa cells droplet encapsulation

HeLa cells were cultured in a Dulbecco’s Modified Eagle Medium (DMEM) (high glucose, pyruvate) supplemented with a final concentration of 10% fetal bovine serum and 1× antibiotic-antimycotic (Gibco, Waltham, MA, USA).

When the HeLa cells reached ≈80% confluence in the cell culture dish, the medium was carefully aspirated using a Pasteur pipette. Subsequently, 5 mL of 1× PBS (137 mm NaCl, 2.7 mm KCl, 10 mm Na2HPO4, and 1.8 mm KH2PO4, pH 7.4) was gently added to the side of the dish and then aspirated. Next, 2 mL of trypsin-EDTA buffer was added, and the dish was shaken by hand to ensure coverage of the bottom. The cells were incubated for 2 min. Next, 8 mL of medium was added, and the total solution (10 mL) was transferred into a 15-mL conical tube and centrifuged at 500 × *g* for 5 min. The resulting cell pellet was resuspended in 1x PBS and washed three additional times by centrifugation at *500 x g* for 5 min. After the final wash, the cells were resuspended in 750 µL of 1 x PBS and used immediately for agarose µ-droplets generation.

### Generation of droplets using microfluidic device

Agarose µ-droplets were generated using a microfluidic droplet generator under controlled flow and temperature conditions. Mineral oil (Sigma-Aldrich, catalog no. M3516) containing 3% SPAN 80 (Sigma-Aldrich, catalog no. 85548) was used as the continuous phase, while the dispersed phase consisted of agarose-based aqueous solutions prepared according to the specific droplet type. Prior to droplet generation, agarose solutions were heated to 65 °C to ensure complete dissolution. *To prevent premature gelatin of agarose within the device, the inlet region of the microfluidic chip was maintained at 42 °C using a flexible insulated heater (OMEGA Engineering Inc, Kapton® (Polyimide Film), insulated heater KHLV Series, 28 V)* connected to a DC power supply for temperature control. The dispersed and continuous phases were then injected into the droplet generator through tubing connected to a syringe pump (NE-1000, New Era Pump Systems Inc., Wantagh, NY), with the syringe diameter set to 5 nm. Flow rates were set to 25 µL/min for the continuous phase and 27 µL/min for the dispersed phase.

For agarose-only control droplets, the dispersed phase consisted of 0.75%(w/v) ultra-low-gelling-temperature agarose (Sigma-Aldrich, catalog no. A5030) dissolved in PBS (137 mM NaCl, 2.7 mM KCl, 10 mM Na_2_HPO_4_, and 1.8 mM KH_2_PO_4_; pH 7.4), and the continuous phase contained 0.75% (w/w) SPAN 80. To generate HeLa cell-containing droplets, a suspension of HeLa cells was mixed with 0.75% (w/v) ULGT agarose prior to droplet formation, and mineral oil supplemented with 3% (w/w) SPAN 80 was used as the continuous phase. For magnetic bead-containing droplets, Ni-NTA magnetic beads (HisPur™ Ni-NTA Magnetic Beads, Thermo Fisher Scientific, catalog no. 88831) were mixed with 1% (w/v) low-gelling-temperature (LGT) agarose (Sigma-Aldrich, catalog no. A4018) before droplet generation, and mineral oil containing 3% (w/w) SPAN 80 served as the continuous phase.

Following droplet generation, the resulting µ-droplets were cooled at 4 °C for more than 3 h to allow complete solidification of the agarose matrix. After agarose solidification, the oil phase was removed by repeated buffer exchange. For agarose-only droplets, magnetic bead-containing droplets, and HeLa cell-containing droplets intended to maintain intact cells, the oil phase was replaced with fresh PBS through multiple washing steps. In contrast HeLa cell-containing droplets designated for cellular permeabilization were subjected to repeated buffer exchange using fresh TE (10 mM Tris-HCl and 1 mM EDTA; pH 8.0) buffer. This post-solidification the desired buffer environment for each droplet type prior to subsequent experimental processing.

### Two-dimensional gel electrophoresis

Cell pellets were washed twice with icecold PBS and sonicated for 10 s by Sonoplus (Bandelin electronic, Germany). Then, they were homogenized by mortor-driven homogenizer ((PowerGen125, Fisher Scientific) in sample lysis solution (7 M urea, 2 M thiourea, 4% (w/v) 3-[(3-cholamidopropy) dimethyammonio]-1-propanesulfonate (CHAPS), 1% (w/v) dithiothreitol (DTT) and 2% (v/v) pharmalyte and 1 mM benzamidine). Proteins were extracted for 1 h at 25 °C with vortexing. After centrifugation at 15000 g for 1 h at 15 °C, insoluble material was discarded, and soluble fraction was used for twodimensional gel electrophoresis. IPG dry strips were equilibrated for 12-16 h with sample lysis solution and respectively loaded with 300 µg of sample. Isoelectric focusing (IEF) was performed at 20 °C using a Multiphor II electrophoresis unit and EPS 3500 XL power supply (Amersham Biosciences). For IEF, the voltage was linearly increased from 150 to 3500 V during 3 h for sample entry followed by constant 3500V, with focusing complete after 96 kVh. Before the second dimension, strips were incubated for 10 min in equilibration buffer (50 mM Tris-Cl, 6 M urea, 2% SDS, 30% glycerol, pH 6.8), first with 1% DTT and second with 2.5% iodoacetamide. Equilibrated strips were inserted into SDS-PAGE gels. 2D gels were run at 20 °C for 1700 Vh. And then 2D gels were Alkaline Silver stained without fixing and sensitization step.

### Expression and purification of BRD4 BD1

pHis-BRD4 BD1 plasmid (Addgene plasmid #196544), originally deposited by Byung II Lee, was obtained from Addgene (Watertown, MA, USA). The constructed BRD4 BD1 plasmid was transformed into E. coli BL21 (DE3). The BL21 cells harboring the BRD4 BD1 plasmid were incubated in Luria Broth (LB) medium supplemented with kanamycin at 37 °C for 16 h overnight. Next, 1 mL of the overnight culture was transferred to 100 mL of LB medium containing kanamycin and incubated at 37 °C until the optical density at 600 nm (OD600) reached 0.4–0.6. Protein expression was induced by the addition of 1 mM IPTG, followed by incubation for 16 h overnight at 20–25 °C with shaking at 200 rpm. After cell lysis by ultrasonication for 15 min and centrifugation at 10,000 rpm (12,298 × g) for 10 min, the clarified lysate was subjected to affinity chromatography using Ni–NTA agarose resin. His-tagged BRD4 BD1 protein was eluted with a buffer containing 50 mM Na2HPO4, 300 mM NaCl, and 250 mM imidazole (pH 8.0). The purified protein was buffer-exchanged into phosphate-buffered saline (PBS; 137 mM NaCl, 2.7 mM KCl, 10 mM Na2HPO4, and 1.8 mM KH2PO4, pH 7.4) and stored at −20 °C with the addition of glycerol [35].

### Synthesis of JQ1-oligo-Cy5 conjugate

Cy5-labeled and thiol-modified oligonucleotides were purchased from Bioneer (Daejeon, Korea). The oligonucleotides were dissolved in 1x TE buffer (10mM Tris-HCl, 1mM EDTA, pH 8.0) to a final concentration of 800 µM. Tris(2-carboxyethyl)phosphine hydrochloride (TCEP; Sigma-Aldrich, catalog no. C4706) was prepared at 8mM in HEPES buffer (50mM, pH 7.0), which was obtained by diluting a 1M HEPES stock solution (Sigma-Aldrich, catalog no. 83264). For reduction of thiol groups, 10 µL of the oligonucleotide solution was mixed with 100 µL of the TCEP solution and incubated at room temperature for 3 h.

Following thiol reduction, excess TCEP was removed by ethanol precipitation. 3M Sodium acetate (Sigma-Aldrich, catalog no. S2889) 11 µL and 100% ethanol 220 µL were added to the reaction mixture, followed by incubation at −80 °C for 1 h. The sample was centrifuged at 13,000 rpm for 30 min at 4 °C to pellet the oligonucleotide, after which the supernatant was carefully removed. The resulting pellet was then washed once with 70% ethanol and centrifuged again at 13,000 rpm for 10 min at 4 °C. Residual ethanol was removed under a gentle stream of argon gas, and the pellet was briefly airdried prior to subsequent processing. Recovered oligonucleotide was resuspended in 190 µL HEPES buffer (50mM, pH 7.0).

Reduced oligonucleotides were reacted with 16 µL of 10 mM (+)-JQ1 maleimide (Tocris Bioscience, catalog no. 7576), and the reaction mixture was incubated overnight at room temperature. The resulting conjugate was subsequently purified by reverse-phase high-performance liquid chromatography (RP-HPLC), and the appropriate fraction were collected, lyophilized, and resuspended in ACN and TE.

### Fluorescence Microscopy

The microscope system comprised an inverted microscope (Olympus IX70, Tokyo, Japan) equipped with a 10× Olympus UPlanSApo objective lens and illuminated LED light source (SOLA SM II light engine, Lumencor, Beaverton, OR). The light passed through the corresponding filter sets (Semrock, Rochester, NY) to excite the fluorescent dye. Fluorescence images were captured using a scientific-grade complementary metal-oxide-semiconductor digital camera (2048 × 2048, Prime sCMOS Camera, Photometrics, Tucson, AZ) and stored in 16-bit TIFF format using Micro-manager software. ImageJ was employed for image processing.

### BRD4 overexpressing HeLa cell preparation

The GFP-BRD4 expression plasmid (Addgene plasmid #65378) was originally obtained from Addgene and kindly provided by a collaborating laboratory. HeLa cells were transiently transfected with GFP-BRD4 expression plasmid using Lipofectamine™ 3000 Transfection Reagent (Thermo Fisher Scientific, catalog no. L3000001) according to the manufacturer’s protocol. A total of 10 µg of plasmid DNA was used per transfection, and cells were harvested 36 h post-transfection for subsequent experiments.

### Cell culture, transfection and Sample Preparation for DNA-PAINT

COS-7 Cells were cultured in high-glucose Dulbecco’s modified Eagle’s medium (DMEM) supplemented with 10% fetal bovine serum (FBS) at 37 °C in a humidified atmosphere containing 5% CO_2_. The cells were electroporated (MPK5000, Invitrogen, CA, United States) during the general subculture, with ∼500 ng of proper plasmid. After 24–48 h of transfection, the cells were fixed in 4% for 10 min at room temperature. For BRD4 labeling, the samples were incubated with 100nM of JQ1-conjugated BP-DNA docking strand 5xR2 (sequence: 5’-TTACCACCACCACCACCACCAGACCACATCGATTTGGGAGTCA-JQ1-3’) for 1 hour at room temperature. For two-color imaging with BRD4-GFP, an anti-GFP nanobody conjugated with 5xR6 docking strand (sequence: 5’-AACAACAACAACAACAACAA-3’, Massive Photonics) was simultaneously applied at a 1:50 dilution.

### Super-Resolution Imaging Protocol and localization cluster analysis

All super-resolution imaging was performed on a commercial spinning disk confocal system (Dragonfly 600, Andor). Coverslips were mounted in a magnetic chamber (Chamilde, LCI) and fitted with tubing for two-color imaging. DNA-PAINT imaging was obtained by illuminating the samples with a high-power 561 nm laser for 10,000 frames at 80−100 ms exposure, using 2 nM dual labeled Cy3b imager R2 (sequence: 5’-Cy3b-TGGTGGT-Cy3b-3’, biomers) and 4 nM Cy3b imager R6 (sequence: 5’-TGTTGTT-Cy3b-3’, massive photonics)in Buffer C (500 mM NaCl in PBS). Raw data from the DNA-PAINT experiments were processed using the Picasso software suite. Single-molecule localization coordinates were extracted using the localize module, and super-resolution images were reconstructed using the render module. Furthermore, the SMLM clusterer function was employed to identify and specify the positions of individual BRD4 and JQ1 molecules by collecting clusters containing at least 10 localizations (k>10) within a 10nm radius.

### DEL screening

For large-scale DEL screening, four conditions were used: (1) magnetic bead-containing droplets, (2) BRD4-bound magnetic bead-containing droplets, (3) permeabilized HeLa cell-containing droplets, and (4) BRD4-overexpressing HeLa cell-containing droplets. For small-scale DEL screening experiments, agarose-only droplets were additionally included as a control condition. For all screening conditions, reactions were performed in micro-centrifuge tubes with a total volume of 250 uL, of which 100 uL corresponded to the droplet phase.

For DEL screening, a screening buffer was used for library incubation, and wash buffers were used for post-incubation washing steps. The screening buffer consisted of PBS supplemented with 0.2 mg/mL Deoxyribonucleic acid, low molecular weight from salmon sperm (Sigma-Aldrich, catalog no. 31149), while the wash buffer was prepared by adding 0.05% Tween-20 to the screening buffer.

For magnetic bead-containing droplet screening, 250 pmol of BRD4 was incubated with magnetic bead-containing droplets at room temperature for 1 h to allow protein immobilization. The beads were then washed three times with screening buffer (1 mL per wash, 3 min each). For HeLa cell droplet-based screening, the droplets were subjected to three sequential buffer exchanges with screening buffer (1 mL per wash, 3 min each).

Two independent DNA-encoded libraries (DELs) were incubated in parallel: (i) a DEL synthesized at the Korea Research Institute of Chemical Technology (KRICT), and (ii) a DEL synthesized at Korea University. For small-scale DEL screening, the library (0.7 pmol/µL) was diluted 1:10 in screening buffer, and 1 µL of the diluted DEL (0.07 pmol) was added to each sample. For large-scale DEL screening, three core libraries were prepared at 1.78 ng/µL, and 10 µL of each core library was combined to generate a total volume of 30 µL. This mixture was then aliquoted at 7.5 µL per sample for four screening conditions. Each DEL was incubated with the respective droplet samples for 2 h at room temperature on a rocker.

Following incubation, magnetic bead-containing droplets were washed six times with wash buffer (1 mL per wash, 3 min each), and HeLa cell-containing droplets were washed six times with wash buffer (750 uL per wash, 3 min each) to remove unbound library members. After washing, magnetic bead-containing droplets were incubated with 50 uL of elution buffer (50 mM Na2HPO4, 300 mM NaCl, and 250 mM imidazole, pH 8.0) to disrupt Ni–NTA/His-tag interaction, thereby releasing BRD4 and associated DEL DNA into the supernatant. The supernatant obtained from magnetic bead-containing droplets was used as the template for subsequent PCR amplification. For HeLa cell-containing droplets, washed droplets themselves were directly used as the PCR template.

For large-scale DEL screening, recovered DEL DNA from all four conditions was amplified by PCR using AccuPower® ProFi Taq PCR PreMix (Bioneer, catalog no. K-2631) with library-specific primers according to the manufacturer’s instructions. PCR was performed under the following cycling conditions: initial denaturation at 95 °C for 3 min; 30 cycles of 95 °C for 20 s, 48 °C for 20 s, and 72 °C for 30 s; followed by a final extension at 72 °C for 10 min. PCR products were purified using the AccuPrep® PCR/Gel Purification Kit (Bioneer, catalog no. K-3038) following the manufacturer’s protocol for PCR purification, and subjected to downstream sequencing analysis.

For small-scale DEL screening, different downstream analyses were applied depending on the experimental condition. For magnetic bead–containing droplets (conditions 1 and 2), eluted samples were amplified by PCR using ProFi Premix and purified prior to analysis. For permeabilized and BRD4-overexpressing HeLa cell–containing droplets as well as agarose-only control droplets (conditions 3–5), washed droplets were directly subjected to quantitative PCR (qPCR) analysis using AccuPower® 2X GreenStar™ qPCR MasterMix (−ROX Dye) (Bioneer, catalog no. K-6254), without an intermediate purification step. qPCR was performed with an initial denaturation at 95 °C for 5 min, followed by 35 amplification cycles consisting of 93 °C for 20 s, 50 °C for 20 s, and 72 °C for 15 s, and a final extension at 72 °C for 10 min. Fluorescence signals were collected during the 72 °C extension step of each cycle.

### Nanopore sequencing

Following PCR amplification and purification, DNA concentrations were measured and normalized to equal DNA mass. Normalized samples were then pooled and subjected to ligation-based nanopore library preparation using a ligation sequencing kit (SQK-LSK 114) following a protocol from community.nanoporetech.com. In brief, each library was subjected to DNA repair and end-prep by a NEBNext FFPE DNA Repair Mix and Ultra II End-prep Enzyme mix. After a purification step using AMPure XP magnetic beads, the sequencing adapters were ligated to both ends of the libraries by NEBNext Quick T4 DNA Ligase and Ligation Adaoter (LA). After a final product clean-up using the Short Fragment Buffer (SFB), each sequencing library was loaded into an R10.4.1 version flow cell (FLO-MIN114, ONT). DNA sequencing was performed on a MinION sequencer (MK1B, ONT). Basecalling was performed using MinKNOW software (Oxford Nanopore Technologies) with a minimum quality score threshold of Q10. Sequencing reads were obtained in FASTQ format. Subsequent data processing and analysis, including read counting and enrichment analysis, were carried out in R using RStudio (R Foundation for Statistical Computing) [36].

